# Sampling the Zebrafish gut Microbiota – A Genome Resolved Metagenomic Approach

**DOI:** 10.1101/2023.12.11.571059

**Authors:** Eiríkur Andri Thormar, Søren Blikdal Hansen, Louise von Gersdorff Jørgensen, Morten T. Limborg

## Abstract

The zebrafish is a promising model organism in the field of functional microbiota research. However, studies on the functional landscape of the zebrafish gut microbiota through shotgun based metagenomics are scarce. Thus, there is a lack of consensus regarding an appropriate sampling method that accurately represents the zebrafish gut microbiota. To address this, we sought to systematically test and evaluate four different methods of sampling the zebrafish gut microbiota: collection of feces from the tank, the whole gut, intestinal content and the application of ventral pressure to facilitate extrusion of gut material. In addition, we included water samples as an environmental control to address the potential influence of the environmental microbiota on the data interpretation. To compare these sampling methods in a context of microbiota-based studies we employed a combination of genome resolved metagenomics and 16S metabarcoding techniques. We observed differences among sample types on all levels including sampling, bioinformatic processing, metagenome co-assemblies, generation of metagenome-assembled genomes (MAGs), functional potential, MAG coverage and micro-diversity. Furthermore, our comparison to the environmental control demonstrated the potential impact of the environmental contamination on data interpretation. The findings emphasise the importance of considering the choice of sampling method. While all sample types tested are informative about the zebrafish gut microbiota, the optimal sample type depends on the specific objectives of the study.

## INTRODUCTION

Zebrafish (*Danio rerio*) is one of the most studied model organisms to date. The use of the model dates back to the research of George Streisinger in the late 1960’s and is today used in a wide variety of research topics from evolutionary biology to human health studies (Grunwald & Eisen, 2002). Todays extensive use of the model is well grounded in that zebrafish are relatively easy to rear, they are relatively cheap to care for, they are easy to breed, they reach adulthood quickly, they have a well defined embryology (Spence et al., 2008), and they are easy to genetically modify (Hwang et al., 2013; Tennant et al., 2019), not to mention the plethora of existing zebrafish related scientific literature – over 53.000 publications to date (*PubMed Search Results (2023)*). It is no surprise then that interest in applying zebrafish as a model organism to study the increasingly popular topic of host-microbiota interactions is on the rise, and a plethora of research papers, preprints and reviews have been published on the matter, ranging from probiotic effect on behaviour (D. J. Davis et al., 2016; Stagaman et al., 2020) to effects of the immune-cell-development gene, *irf8*, on the zebrafish microbiota (Earley et al., 2018). Expectedly it seems that the most prevalent method of studying the zebrafish gut microbiota is using 16S metabarcoding, a valuable and powerful tool to investigate and describe the microbiota composition, diversity and taxonomic landscape of the gut microbiota. However, given the increasing interest in the zebrafish model for studying host-microbiota interactions and advances in development of sequencing technologies and bioinformatic tools, it is interesting to consider the number of studies applying more functional approaches. To date, three publications (Kayani et al., 2021, 2022; Zhang et al., 2021) and a single preprint (Gaulke et al., 2020) appear to form the entirety of the zebrafish short read based metagenomics literature. Only one of which (Kayani et al., 2022) uses a genome resolved metagenomics approach, that is de novo assembly of whole bacterial genomes often termed metagenome assembled genomes (MAGs). One of the studies relies on sampling whole intestines (Zhang et al., 2021), while the others all rely on the same sampling methodology to characterise the gut metagenome – fecal matter sampled directly from the tank.

This leads to considerations regarding sampling strategy, i.e what type of sample to take that best represents the zebrafish intestinal microbiota for a metagenomic shotgun sequencing study while providing enough microbial DNA reads. For sampling the zebrafish microbiota there are several different methods noted in the literature, (Stagaman et al., 2020) but little tangible consensus on what constitutes an “appropriate” sample. These sampling methods all differ in degree of invasiveness and relative ease of sampling. The first and seemingly most prevalent one is the collection of fecal matter from the water, an easy sample to acquire and completely non-invasive; such sampling even allows for a time series study to be conducted. However, this approach provides a tank-level representation of the zebrafish microbiota, lacking individual fish representation. This problem can however be overcome as done in (Gaulke et al., 2019) where individual fish are separated into smaller tanks and fecal matter subsequently collected, potentially a rather time consuming effort but effective. Even though this problem with individual variation is overcome another less obvious problem arises – the water microbiota. As the sample is acquired directly from the tank, the boundary between the environment and water may be blurred, potentially rendering functional and taxonomic insights less than optimal. A second way of sampling the microbiota is to dissect out the whole gut. This is an invasive and rather time-consuming technique but does carry the benefit of individual association and no direct contact with the surrounding water. The main drawback with this method is that most of the DNA extracted will likely be host associated DNA, a common feature of fish microbiota studies (Collins et al., 2021; Hennersdorf et al., 2016; Rasmussen et al., 2021; Riiser et al., 2019), and something we have previously observed in whole guts of zebrafish (unpublished data). A high amount of host DNA can impact the quality and resolution of the analysis of the microbiota (Pereira-Marques et al., 2019). A third possible method would be to dissect out the entire gut and dissect out the intestinal content, as done in (Gaulke et al., 2016). Although this method is rather time-consuming and invasive it has the potential to get both individual level variation and potentially more microbial DNA. A fourth potential option to consider would be to press along the underside from the head towards the urogenital tract and extract the emerging intestinal content. There are many more potential sampling techniques that may cover the nieces of the zebrafish gut microbiota i.e. differential composition and function in different sections of the gut. We will however only focus on four different sampling techniques that each should represent the whole zebrafish gut microbiota.

We sought to address the sampling of the zebrafish intestinal microbiota by comparing four different methods throughout a genome resolved metagenomic pipeline. We compared feces sampled directly from the tank (here” feces”, whole gastrointestinal tract samples (here “whole gut”), pressure on the underside (here termed “squeezed gut”), and finally intestinal content dissected from whole guts (here termed “intestinal content”). Water samples were included as an environmental control and for estimation of environmental contamination. Differences among sample types were evident on all levels from sampling to analyses. We conclude that the method of choice should depend largely upon the aim of the study and that special care has to be taken concerning environmental contamination.

## METHODS

### Zebrafish husbandry and ethics statement

The zebrafish were reared in a re-circulatory system (Techniplast active blue) with a dark/light cycle of 14/10 h. The zebrafish were fed with dry feed (ZM-300, ZM Fish food and Equipment, UK) and live Brine Shrimp (*Artemia* sp., ZM Fish food and Equipment, UK). The zebrafish collected for this study were all housed in the same tank. The experiment was conducted in accordance with a permit from The Animal Experiments Inspectorate under the Danish Ministry of Environment and Food (Permit: 2021-15-0201-00951). During sedation and sacrifice of fish tricaine methanesulphonate (MS222, A5040, Sigma-Aldrich, Denmark) was used.

### Sampling

As described above, we aimed to compare four different approaches for their efficiency in sampling DNA for characterising the zebrafish gut metagenome using shotgun sequencing. First, fecal matter was sampled directly from the tank using a single-use polyethylene pipette and transferred to a petri dish. Using an inoculation loop, approximately ½ of a 10 ul loop, fecal matter was then transferred to and stored in an MP Biomedicals Lysing E Matrix tube with 0.5 ml of 1X DNA/RNA Shield (Zymo research). A total of three replicate fecal samples were collected.

Second, the whole gastrointestinal tract of zebrafish was then carefully dissected out and stored in an MP Biomedicals Lysing E Matrix tube with 0.5 ml of 1X DNA/RNA Shield (Zymo research). A total of three whole gastrointestinal tracts were collected.

Third, we sampled intestinal content from the zebrafish by gently “squeezing”/putting pressure along the abdominal side of the fish from the pectoral fin towards the anal fin. We sampled the resulting excretion from the urogenital pore using an inoculation loop, approximately ½ of a 10 ul loop fecal matter was then transferred to and stored in an MP Biomedicals Lysing E Matrix tube with 0.5 ml of 1X DNA/RNA Shield (Zymo research). A total of three “squeezed gut” samples were collected.

Fourth, the gastrointestinal tract was carefully dissected out and the tip of a single-use inoculation loop was gently traced along the entire dissected gastrointestinal tract to extrude the intestinal contents avoiding tearing of the intestinal tract. Using an inoculation loop, approximately ½ of a 10 ul loop of the intestinal content was then transferred to an MP Biomedicals Lysing E Matrix tube with 0.5 ml of 1X DNA/RNA Shield (Zymo research). Four intestinal-content samples were collected.

Finally, to serve as the environmental control, 500 ml water was collected 50 ml at a time and filtered using a sterivex filter. The filter was then filled with DNA/RNA Shield (Zymo research) before storage. All samples were stored at –20 °C before DNA extraction.

### DNA extraction and sequencing of metagenomic and 16S data

Prior to DNA extraction all samples were lysed in a Tissuelyzer (Qiagen) at 30 GHz for 5 minutes and spun down at 13.000 rpm for 5 min. Subsequently 400 ul lysate from each sample was transferred to a 96 deep well plate in a randomised order. Three DNA/RNA shield (Zymo research) negative extraction controls were included in the process. DNA was extracted following the manufacturer’s recommendations of the magnetic bead based Quick-DNA MagBead Plus Kit (Zymo Research). The DNA concentration of each sample was measured using a Qubit fluorometer (Invitrogen). A part of the eluted DNA was shipped to Novogene (Cambridge, UK) for metagenomic shotgun sequencing where DNA was randomly sheared into short fragments by sonication. Library was prepared using Novogene NGS DNA Library Prep Set (Cat No. PT004). The library was quantified with a Qubit fluorometer and real-time PCR and bioanalyzer 2100 (Agilent) was used for size distribution detection. The quantified libraries were then pooled and subject to 150 paired-end sequencing strategy on a NovaSeq 6000 instrument (illumina).

The leftover DNA was used for generating paired 16S data. First the V3-V4 hypervariable region of the 16S rRNA gene was amplified using 341F (5’-CCTAYGGGRBGCASCAG-3’) and 806R (5’-GGACTACNNGGGTATCTAAT-3) (Yu et al., 2005) with the addition of 20 different tags to the 5’ end of each primer. A negative PCR control was also included Each reaction was 25 ul and consisted of 13.5 ul ddH20, 2.5 ul 10X TaqGold buffer (GeneAmp®), 2.5 ul Mgcl2(25mM), 1.5 ul BSA (20ng/ul), 0.5 ul dNTPs(10 mM), 0.5 ul DNA polymerase AmpliTaq Gold® (5 U/ul), 2 ul primer mix and 2 ul DNA extract (2 ul ddH20 for the PCR control). The PCR conditions were as follows: Initial denaturation at 95°C for 10 min followed by 30 cycles of denaturation at 95 °C for 15 sec, annealing at 53 °C for 20 sec and extension at 72 °C for 40 sec followed by a final extension at 72 °C for 10 min. The amplified PCR products were then pooled in an equimolar fashion based on agarose gel band brightness and purified using the HighPrep PCR Clean-up System® (MagBio Genomics Inc). Subsequently the PCR-free single-tube metabarcoding library preparation protocol, Tagsteady (Carøe & Bohmann, 2020), was used to construct the library. The library was quantified using the NEBNext® Library Quant Kit for Illumina® (New England Biolabs) by mixing 2 ul 1:10.000 diluted library with 8 ul Quant mastermix with added primers (New England Biolabs). Due to low initial concentration of adapter ligated product we made triplicates of the library, pooled the triplicates and upconcentrated using a vacuum concentrator, Concentrator plus® (Eppendorf). The library was then sequenced at the GeoGenetics Sequencing Core, University of Copenhagen, Globe institute using an Illumina MiSeq platform, reagent kit v3 at 600 cycles. Negative extraction controls and the PCR control were included in the library and sequenced.

### Bioinformatic processing

Quality of raw reads was assessed using FastQC (Andrews, 2010) and MultiQC (Ewels et al., 2016) for subsequent Quality filtering steps. AdapterRemoval/v.2.3.3 (Schubert et al., 2016) was used for removal of low quality reads and adapters. Duplicates were removed using the rmdup function in Seqkit/v.2.3.1 (Shen et al., 2016) and reads re-paired using bbmap/v.38.84 (Bushnell, 2014). Host reads were filtered out by mapping the quality filtered reads to the host genome (GCA_000002035.4) and the Human genome (hg38) with minimap2/2.24 (H. Li, 2018, 2021) using default parameters for short accurate genomic reads, and all unmapped reads were kept. The mapped reads were kept in separate files to assess depth of coverage using the samtools-depth function (Danecek et al., 2021). The metagenomic reads were co-assembled on sample type basis (that is, 5 co-assemblies) using MEGAHIT/v.1.2.9 (D. Li et al., 2015) with metagenomic sensitive presets and a minimum contig length of 1000bp. Quality of assemblies was assessed using Quast/v.5.0.2 (Mikheenko et al., 2018). Further analyses of assemblies, and visualisation was performed using the anvio platform (Eren et al., 2021; Murat Eren et al., 2015).

For each of the five co-assemblies the following was done: i) anvío was used to identify open reading frames (ORFs) using Prodigal/v.2.6.3 (Hyatt et al., 2010). ii) HMMER/v.3.3 (Finn et al., 2011) was used to identify sets of singe-copy-core genes (SCGs) of protista, archaeal, and bacterial origin (Lee, 2019). The HMMs of SCGs were used for estimating the number of recoverable bacterial genomes in the assemblies and for completion and redundancy estimates of MAGs. iii) The ORFs were annotated in the anvío platform using functions from NCBÍs Clusters of Orthologous Groups (COGs) (Tatusov et al., 2003), the KOfam HMM database of KEGG orthologs (KOs) (Aramaki et al., 2020; Kanehisa & Goto, 2000) and Pfams (El-Gebali et al., 2019). iv) Kaiju (Menzel et al., 2016) was used to infer taxonomy of genes with NCBÍs non-redundant protein database ‘nŕ and import into the anvío framework as described here (https://merenlab.org/2016/06/18/importing-taxonomy/). v) For prediction of the number of genomes in the assemblies we used the “anvi-display-contigs-stats” function. vi) Reads were then mapped to contigs using minimap2/2.24 (H. Li, 2018, 2021) and samtools (Danecek et al., 2021) and stored as BAM files. Each BAM file was profiled in anvío sung “anvi-profile” for estimation of coverage and detections statistics of each contig and each profile was integrated into a merged profile database for further processing using “anvi-merge”. vii) Finally, Binning was performed using the anvi-cluster-contigs which uses CONCOCT/v.1.1.0 (Alneberg et al., 2014). Each bin was then manually refined based on tetranucleotide frequency and differential coverage across samples using “anvi-refine”. MAGs were called using “anvi-rename-bins” where each bin that was more than 50% complete and less than 10% redundant was defined as a MAG. Anvío was also used to infer MAG taxonomy based on single-copy core gene sequences from the Genome Taxonomy Database (GTDB) (Parks et al., 2018).

To create a non redundant set of MAGs representative of all samples, sequences for each MAG from each merged profile database were extracted. CheckM (Parks et al., 2015) was used as an additional quality assessment of the MAGs. We then applied dRep (Olm et al., 2017) to dereplicate the set of MAGs into a set of non-redundant species-level representatives of the collection based on average nucleotide identity (ANI) and quality of MAGs based on the previous CheckM quality assessment (Supplementary file S1). We then mapped the metagenomic reads from all samples to the set of non-redundant MAGs, and followed the same steps as detailed above using the anvío platform. For further comparison with existing data we mapped our metagenomic reads to the Zebrafish Fecal v1.0 MAG catalogue in the MGnify genomes resource (Gurbich et al., 2023).

Quality of raw reads from 16S V3-V4 sequencing was assessed using FastQC (Andrews, 2010) and MultiQC (Ewels et al., 2016) for subsequent Quality filtering steps. Cutadapt/v.4.4 (Martin, 2011) was used to trim adapter and primer sequences. Subsequently the DADA2 (Callahan et al., 2016) pipeline was applied for quality filtering and trimming, inferring of sequence variants, merging of forward and reverse reads, removal of chimeric sequences, and taxonomy assignment with the SILVA/v138 reference database (Quast et al., 2013). Then Decontam (N. M. Davis et al., 2018) was applied to remove putative contaminants, after which LULU ((Frøslev et al., 2017) was applied for ASV curation and removal of erroneous sequences. The ASVs were then merged at the genus level for analyses.

### Analyses and visualisation

Rarefaction curves were estimated using the R package *vegan* (Oksanen et al., 2020). The relative abundances of MAGs were calculated using the mean coverage across the MAG and then normalised using the length of the MAG using TMP normalisation with the R package *ADI-impute(Xu et al., 2021)*. Microbiota composition analyses was performed using the R package *phyloseq (McMurdie & Holmes, 2013)*. Visualisation of data was performed using the R package *ggplot2* (Wickham, 2016) the anvío platform (Eren et al., 2021; Murat Eren et al., 2015). The Icons representing sample types, used in figures, were created using BioRender.com.

## RESULTS

### Sample types differed in all stages of sample processing

As an initial comparison of sample types we first ranked them according to two criteria. Relative sampling difficulty, and consistency (Table 1). Consistency refers to how alike the DNA content and microbiota profiles of the same sample type were. The fecal samples were the easiest to sample followed by squeezed gut content, dissected whole gut and then the dissected intestinal content was most difficult to sample. The fecal samples were also most consistent among samples followed by whole gut, dissected gut and then squeezed gut was least consistent among samples. Three of the intestinal content samples did not yield a great amount of sample, while a single one yielded substantially more compared to the others, the same happened to the squeezed gut samples where a single sample clearly yielded more “clean” fecal matter than did the others. In the rest of the results we choose to also include the water samples as they serve as an environmental control.

**Table 1:**
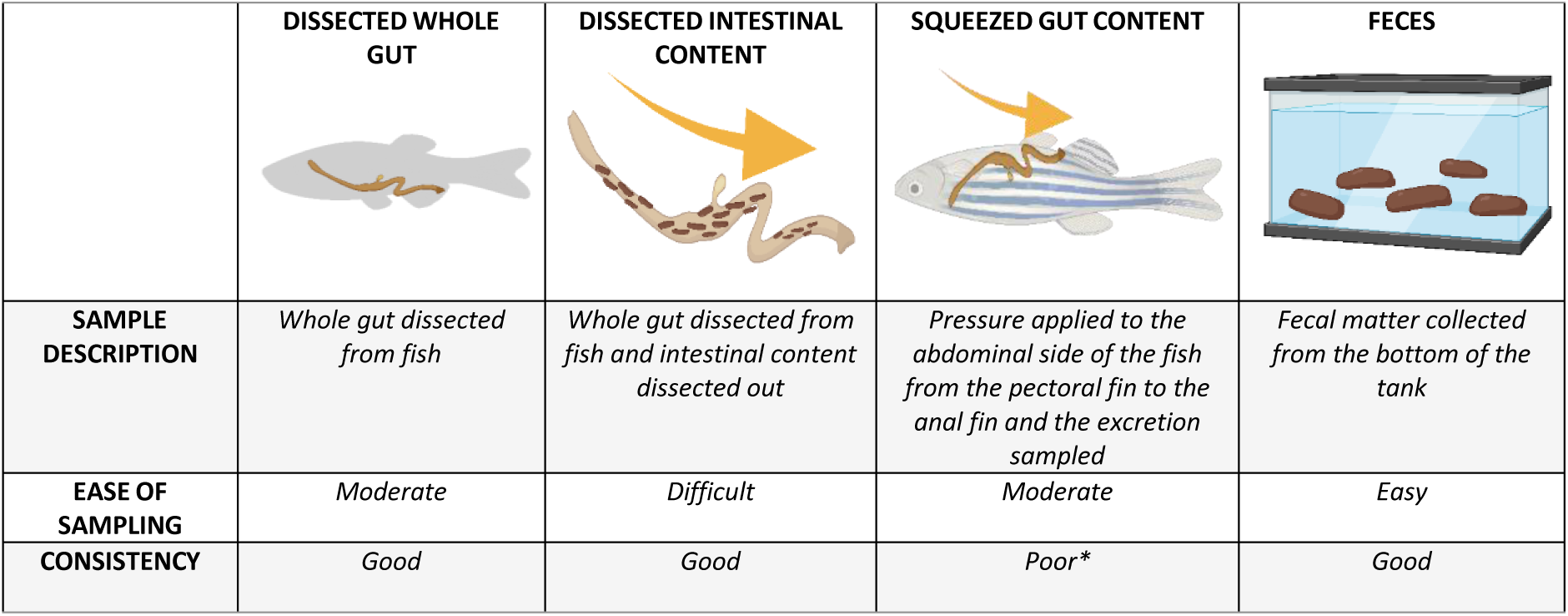
Overview of the sampling process among sample types in terms of feasibility of sampling. *Potential for optimization and improvement in sampling strategy

|  | DISSECTED WHOLE GUT | DISSECTED INTESTINAL CONTENT | SQUEEZED GUT CONTENT | FECES |
| --- | --- | --- | --- | --- |
| **SAMPLE DESCRIPTION** | Whole gut dissected from fish | Whole gut dissected from fish and intestinal content dissected out | Pressure applied to the abdominal side of the fish from the pectoral fin to the anal fin and the excretion sampled | Fecal matter collected from the bottom of the tank |
| **EASE OF SAMPLING** | Moderate | Difficult | Moderate | Easy |
| **CONSISTENCY** | Good | Good | Poor* | Good |

The sample types differed on all levels of sample processing. Here we refer to sample processing as the process of generating MAGs, that is from DNA extraction through bioinformatic processing and metagenomic assembly. After comparing the practicality of sampling we compared the average post-extraction DNA concentration among the different sample types (Fig.1A). As expected, due to the high host DNA content, the whole gut had the highest average DNA concentration, followed by intestinal content and squeezed gut, while feces and water had the lowest average concentration.

**Figure 1:**
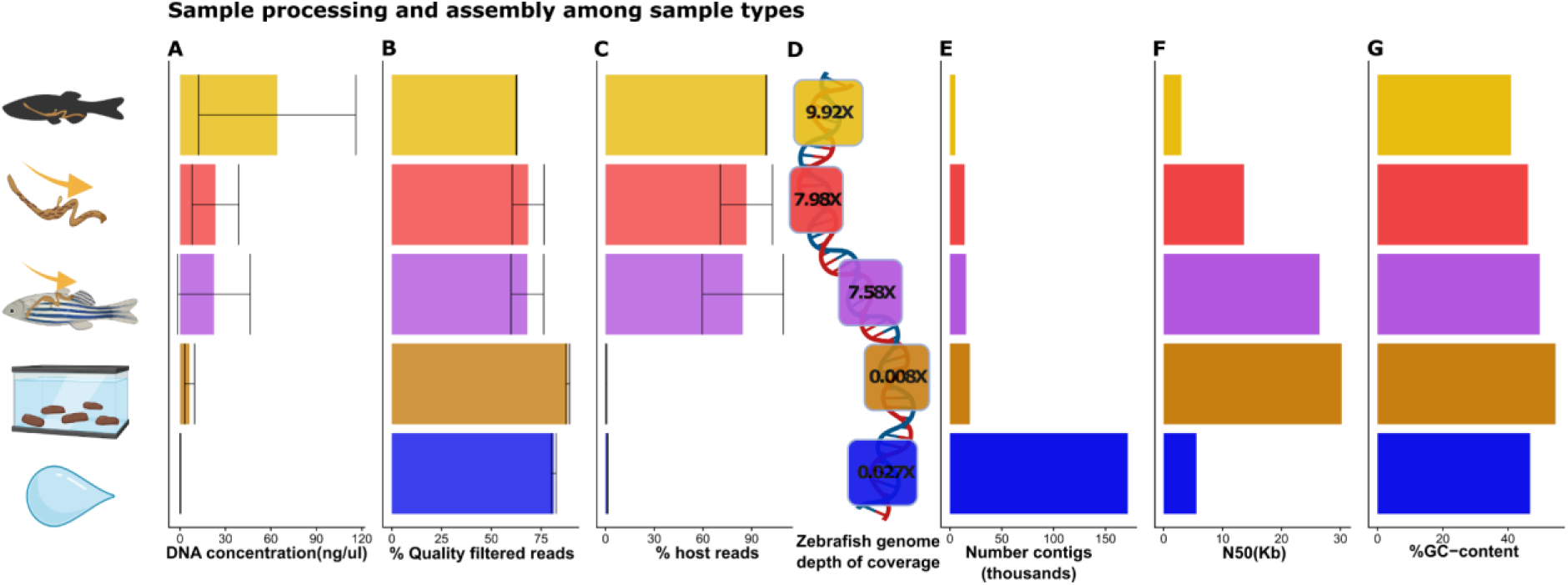
Comparison among sample types through the sample processing. The colours and icons represent different sample types. From bottom to top they represent Whole gut, Intestinal content, Squeezed gut, feces and water A) Mean post extraction DNA concentration among sample types. B) Mean percentage of reads removed after quality filtering and deduplication. C) Mean percentage of quality filtered and deduplicated reads removed after host filtering. D) Mean depth of coverage of the zebrafish reference genome. E) Number of contigs in co-assemblies among sample types F) N50 value of the co-assemblies among sample types, and G) the GC-content of the co-assemblies among sample types. Icons were created using Biorender.com

Metagenomic shotgun sequencing yielded approximately 2.4 billion reads across 15 samples, a single water sample failed sequencing due to low DNA content along with a negative control (indicating the efficacy of the negative control) and were therefore not included in the analysis. After quality filtering and removal of duplicates approximately 647.5 million reads remained. The average percentage of reads removed after quality filtering and removal of duplicates was then compared among sample types (Fig.1B). Whole gut samples had the lowest percentage of removed reads followed by intestinal content, squeezed gut, water and finally most reads were removed from the fecal samples. The standard deviations should also give an indication of the consistency within the sample type group making the intestinal content and squeezed gut less consistent than whole gut and fecal samples. After human and zebrafish filtering approximately 157 million reads remained. The percentage of quality filtered reads that were removed by host filtering was then compared among the sample types to give an indication of the amount of host content among sample types (Fig.1C). As expected the highest amount removed by the host filter was whole gut samples with a strikingly high percentage (99%), followed by intestinal content (86.7%), squeezed gut (84.5%), water (1.29%) and interestingly, the lowest percentage was removed from the feces samples (0.473%). Notably the sheer number of reads not mapping to the host zebrafish genome was highest in the water samples followed by feces, intestinal content, squeezed gut and finally whole gut. The data therefore counterintuitively indicates that a lower initial post-extraction DNA concentration will yield more metagenomic reads in the case of zebrafish shotgun metagenomic sequencing.

Despite this, recovering host genetic data may also be a valuable resource for some studies and for avoiding waste of sequencing depth and cost. We therefore included a comparison of the average depth of coverage of the zebrafish genome recovered from each sample type (Fig.1D). Surprisingly fecal samples presented an unexpectedly lower coverage compared to the water samples. Squeezed gut and intestinal content exhibited relatively similar depths of coverage. As anticipated the whole gut samples had the highest depth of coverage.

Metagenomic assemblies of the unmapped reads were made for all sample types and compared (Fig.1E-G). The assemblies differed at all levels, the number of contigs was by far the highest in the water samples then followed by feces, then the squeezed gut, Intestinal content and finally the lowest number of contigs was in the assembly of the whole gut. As depicted (Fig.1F-G) the GC content and N50 value varied widely among the assemblies of the different sample types.

### Fecal samples provide the highest number and quality of MAGs among sample types

Based on the presence/absence of 71 bacterial single copy core genes (SCGs) the estimated total number of bacterial genomes in the five assemblies was estimated (Fig.2A). The lowest estimated number of genomes was as expected low in the whole gut samples (2 bacterial genomes) followed by, intestinal content (13 genomes), squeezed gut (25 genomes), Feces (24 genomes) and finally water had the highest number of estimated bacterial genomes (49 genomes). After automated binning and manual refinement of the MAGs we compared the number and quality of MAGs. MAGs were called if they were over 50% complete and had less than 10% redundancy, MAGs were classified as high quality if they were over 90% complete and less than 5% redundant. Low quality MAGs less than 50% complete and higher than 10% redundant were not included in the analysis. In total 77 MAGs were called across sample types. The highest number of MAGs (28 MAGs) were called in the water samples (57% of expected MAGs) of those 14 were classified as high quality. A total of 23 MAGs were called from the fecal samples (95% of expected MAGs) of which 12 were classified as high quality.

**Figure 2:**
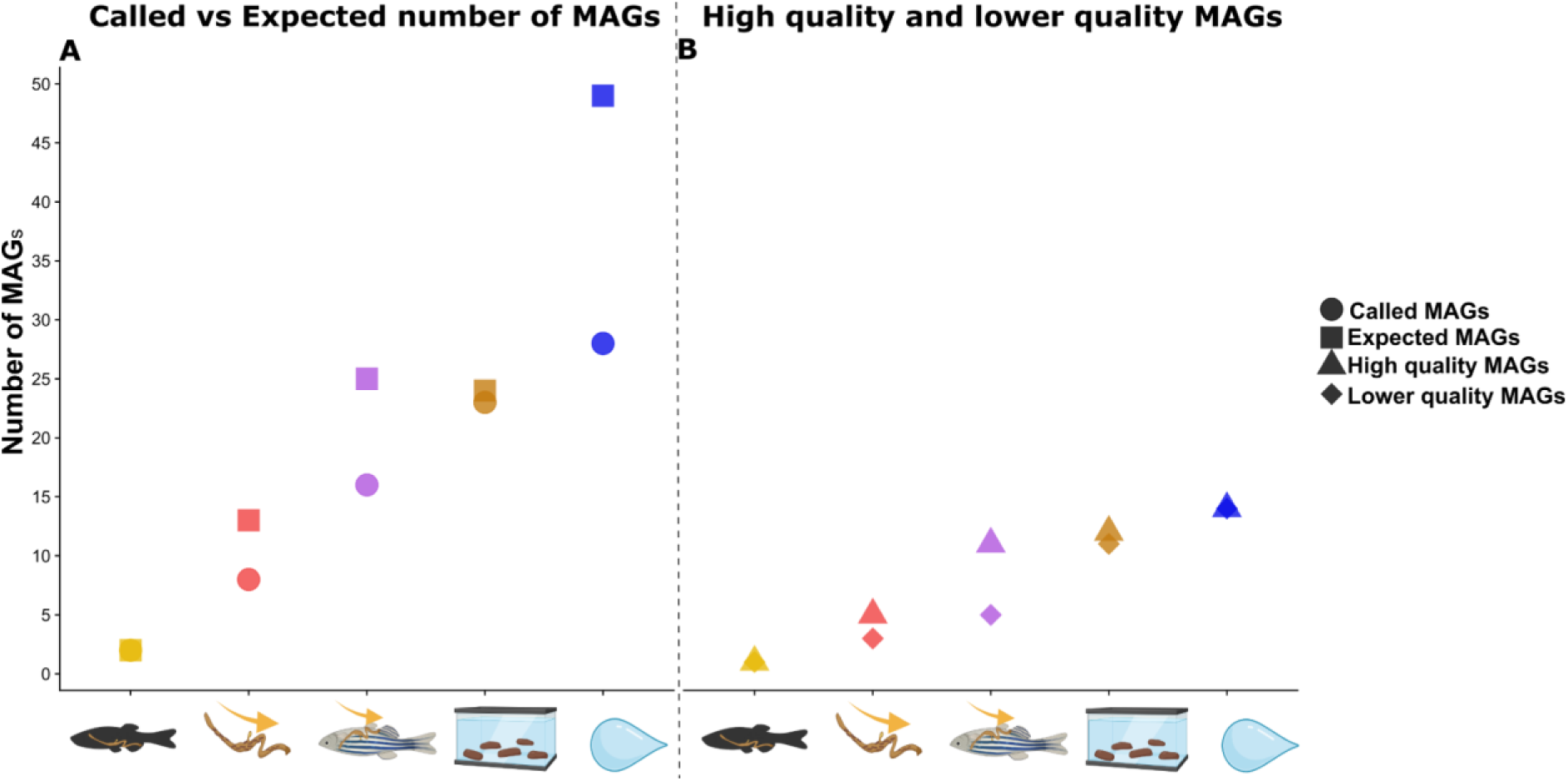
A) plot showing the estimated and actual number of bacterial genomes in the co-assemblies among sample types. The circles represent the called MAGs and the squares represent the estimated MAGs B) plots showing high quality and lower quality MAGs. The triangle represents high quality MAGs (>90% complete and <%5 redundant) while the diamond represents lower quality MAGs (>50% complete and >10% redundant). The colours and icons indicate the sampling type, yellow is whole gut samples, red is Intestinal content samples, purple is squeezed gut samples, brown is feces and blue is water samples.

From the Squeezed gut samples 16 MAGs were called (64% of expected MAGs) 11 of which were classified as high quality. Eight MAGs were called in the intestinal content samples (61% of expected MAGs) of which 5 were classified as high quality. Finally, only two MAGs were called in the whole gut samples, one of which was classified as high quality. It is clear that the representativeness, the number and quality of the assemblies vary widely, so to compare the sample types we created a MAG catalogue consisting of 41 non-redundant MAGs (see supplementary file S1) and mapped the host filtered reads back to it.

First we assessed sequencing depth and generated a rarefaction curve of gene calls across all samples which revealed striking and interesting differences among sample types in terms of data saturation (Fig.3A). Expectedly, as only two MAGs were recovered, the whole gut samples did not reach saturation, indicating insufficient sampling depth. We observed that the intestinal content samples reached saturation indicating sufficient sampling depth. This is in stark contrast to the squeezed gut samples where most of the diversity seems to belong to a single sample while the others do not reach saturation, indicating a lack of consistency in sampling. However, the saturated curve from the single squeezed gut sample is similar to the one of the fecal samples which all saturate around the same time as do the intestinal content samples although with a much higher diversity. The water samples are almost saturated but indicating the highest diversity. This is supported by the coverage of the MAG catalogue among samples.

**Figure 3:**
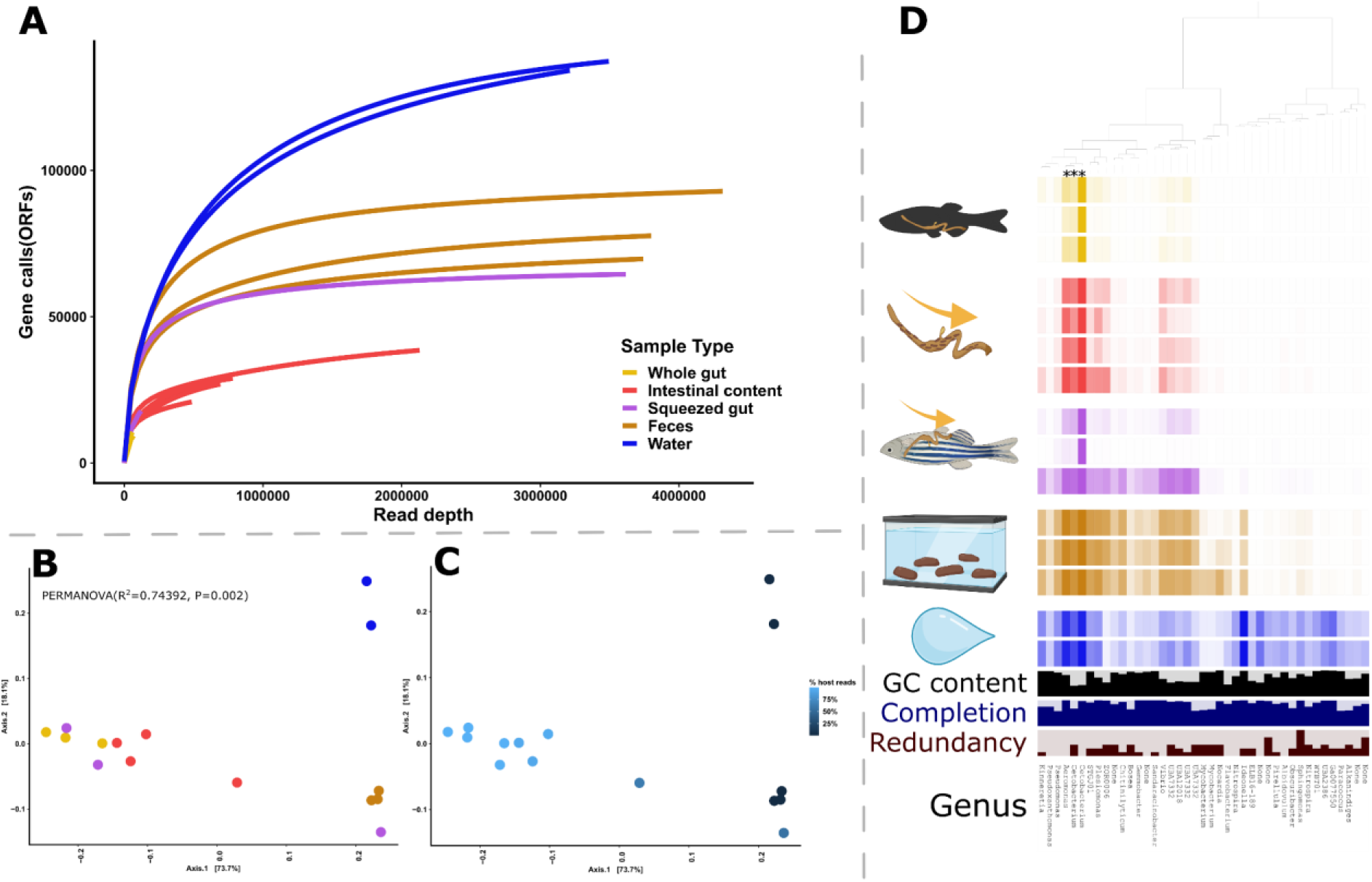
MAG abundance and diversity among sampling types. The colours and icons indicate the sampling type, yellow is whole gut samples, red is Intestinal content samples, purple is squeezed gut samples, brown is feces and blue is water samples. A) Rarefaction curve of ORFs among sample types. B) PCoA of MAGs among samples using Bray-Curtis distance the different colours represent sample type as indicated by the icons. C) PCoA of MAGs where the blue coloured gradient represents the percentage of removed host reads. D) log transformed mean per sample coverage of MAGs in the MAG catalogue created for this study. Each column of square represents a single MAG and each row represents a single sample. GC content of each mag is represented by black bars and completion and redundancy estimates of each MAG is represented by blue and red bars respectively, the genus level of each MAG is also indicated in text. The three MAGs representing over 5% of the overall abundance are indicated with asterixis on the dendogram.

We then looked at the relative abundances of taxa across the dataset (Fig.3D). Of the 41 MAGs only three comprise over 5% of the relative abundance across the sample set, one of which belongs to the genus *Cetobacterium* (assigned species *Cetobacterium sp000799075*) is clearly dominant across all sample types. It comprises the highest percentage in the Whole gut samples (85.2%) followed by Intestinal content (72.1%), Squeezed gut (67.4%), feces (39.8%) and water (38.7%). The second most abundant MAG belongs to the genus *Aeromonas* and its relative abundance is highest in the fecal samples (13.2%), followed by water (12.4%), intestinal content (11.2%), squeezed gut (5.03%) and whole gut (3.62%). The third and final MAG comprising over 5% of the relative abundance also belongs to the Genus Cetobacterium (assigned species *Cetobacterium somerae*). Its relative abundance is highest in the fecal samples (8.38%), followed by intestinal content (7.36%), water (6.19%), squeezed gut (6.156%) and finally whole gut (4.04%). A total of 10 MAGs (the three summarised included) comprise over 1% of the cumulative relative abundance (summary of those can be found in Supplementary file S1). It should be noted that one of the 10 MAGs has by far the highest relative abundance in the water samples, that is a MAG belonging to the genus *Ideonella*. Despite the obvious differences among sample types in terms of relative abundance, significant differences were not observed among the sample types, likely due to the small per-group sample numbers. The beta-diversity of the microbiota composition shows significant difference based on sample type (Fig.3B). The first axis of the PCoA separates the whole gut, intestinal content and part of the squeezed gut content from the fecal and water samples and a single squeezed gut sample (indeed the one that looks more like the fecal samples). The second axis of the PCoA separates the water samples reflecting the different composition of the water compared to the other samples. Moreover, there appears to be a gradient in samples ranging from low-high host DNA content along the first axis of the PCoA (Fig.3C).

Before analysing the functional potential among the sample types we compared the relative abundances of MAGs to 16S amplicon based AVSs generated for the same samples (Fig.4) We focused on comparing the relative abundance of four prevalent taxa ensuring a consistent genus level match between the 16S dataset and the MAG catalogue. These taxa belong to the genera of *Cetobacterium*, *Aeromonas*, *Pseudomonas* and *ZOR0006*. Although not highly significant in all samples, the correlation between log2 transformed relative abundances of the five taxa in 16S and shotgun seems almost perfect among sample types indicating that the functional metagenomic data accurately represents the composition of the microbiota as characterised by the normally used 16S based barcoding approach. The main difference seems to be in the abundances of *Cetobacteriuma* and *Aeromonas* in the fecal and water samples of which there is higher abundance in the 16S data compared to the shotgun data. Diversity and composition analyses were also performed on the 16S data and they largely reflected the composition of the shotgun data although there was less difference between the sample types, i.e. the 16S data from the fecal samples was similar to the whole gut samples (supplementary file S2,Fig.S1).

**Figure 4:**
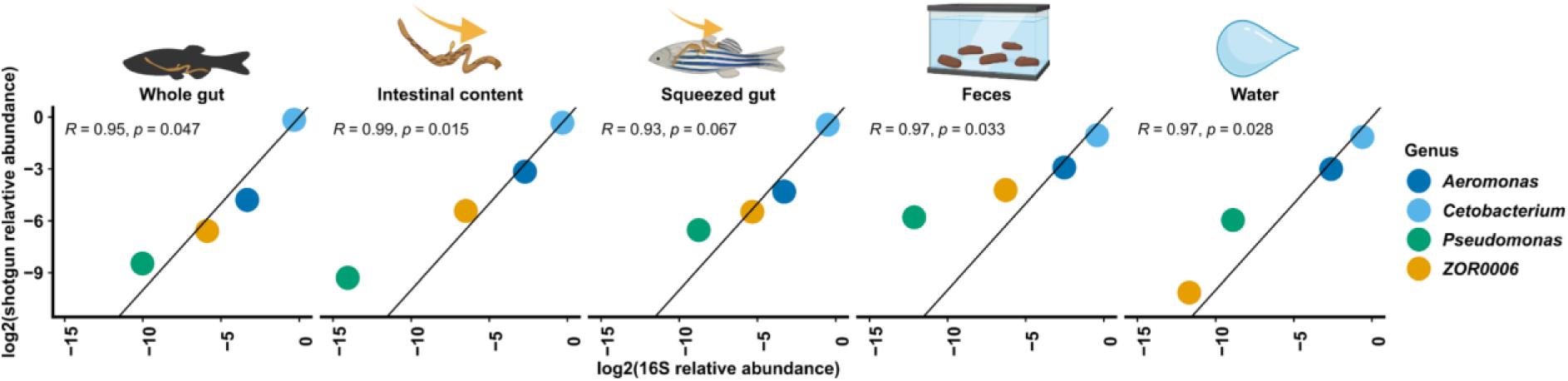
The figure shows the relationship between the log2 transformed relative abundances of the top 4 taxa shared between the 16S metabarcoding data and the shotgun sequencing data among the five sample types. The line represents an x=y relationship. The icons represent the different sampling types and the colours differentiate the four genera.

### Functional composition differs among sample types

Having established the differences in composition among sample types and validation with 16S metabarcoding data we aimed to investigate the differences in the functional composition.

From the difference between the abundance of MAGs among sample types we expect the gene content to vary similarly among the sample types with increasing depth of functional inference (Fig.3). We started by comparing COG categories among sample types (Fig.5). A total of 168.366 genes were called across the MAG catalogue of those 120.410 had assigned COG categories. We compared the sample types in how many gene calls were unique/shared among sample types (Fig.5A) and how much potential functional fraction (that is the fraction of normalised coverage of gene calls) those gene calls contribute (Fig.5B). We identified three main fractions of gene calls among sample types. Firstly, a core fraction of gene calls that are shared among all sample types. Secondly, a fraction shared among all sample types except for whole gut samples was included, likely reflecting the diversity of functions not captured by the whole gut content samples due to high host DNA content. And a fraction shared exclusively between fecal samples and water samples. All sample types shared a core fraction of 35.282 gene calls with assigned COG functions. This fraction of gene calls accounts for most of the potential functional abundance 99.7% (sd=0.0974) in the whole gut samples, 98.8% (sd=0.359) in the intestinal content samples, 95.2% (sd=3.49) in the squeezed gut samples, 90.1% (sd=3.77) in the fecal samples and only 45.1% (sd=7.58) in the water samples suggesting that these functions likely represent a set of core functions in the zebrafish gut microbiota opposed to be sourced from the water microbial community. In the fraction shared among all samples except the whole gut samples (20.967 gene calls) the fraction of functional potential was only 1.02% (sd=0.269) in the intestinal content samples, 4.2 % (sd=3.26) in the squeezed gut samples, 6.79% (sd=1.51) in the fecal samples and 5.16% (sd=0.0203) in the water samples. Thus, likely reflecting the increased resolution of the microbial functions of lower abundance taxa when avoiding sampling of the host gut tissue. Looking at the fraction only shared among the water and the feces (25.736 gene calls) it accounts for only 1.19% (sd=0.728) of the functional potential in the fecal samples in stark contrast with the 33.2% (sd=5.34) in the water samples. Thus likely reflecting a fraction which corresponds to contamination from the water samples. We then removed the fraction of gene calls from the sample set that had a higher mean coverage in the water samples compared to the other sample types. This resulted in a big reduction of shared gene calls both in the fractions shared among only the fecal samples and water samples and in the fraction shared among all sample types apart from whole gut samples. Further addressing this we focused only on the gene calls with COG functions from the 10 MAGs that constituted over 1% of the relative abundance (24.432 gene calls). Subtracting the gene calls from the *Ideonella* MAG, which was highest in abundance in the water samples completely removed the fraction of gene calls (2.604 gene calls) shared between only the fecal and water samples along with a part of the fraction shared between all sample types except the whole gut samples. This analysis was repeated for gene calls with assigned Pfam functions (supplementary material file S2, figure S2) and indicated similar results.

**Figure 5:**
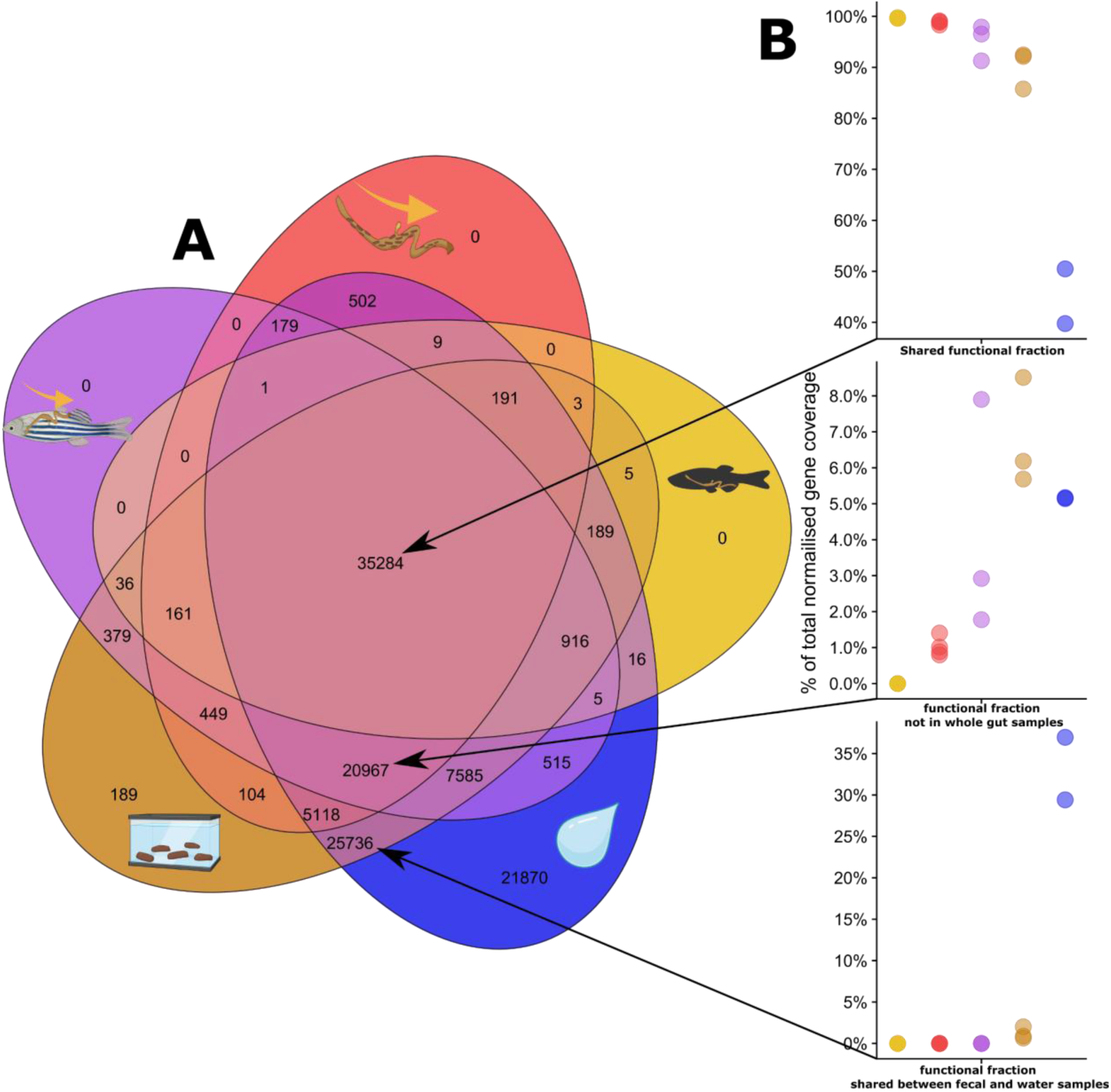
A) Venn diagram of number of gene calls with assigned COG categories across between sample types. B) percentage of normalised gene coverage of genes in fractions of the venn diagram, as indicated by the arrows, among sample types. The colors and icons represent the different sample types. The colours and icons indicate the sampling type, yellow is whole gut samples, red is Intestinal content samples, purple is squeezed gut samples, brown is feces and blue is water samples.

### Variability captured was driven by MAG coverage

To briefly address the increasingly insightful and rising topic of microbial population genetics we profiled the single nucleotide variants (SNVs) variability among sample types (Fig.6). To do this we chose the highest abundant MAG across the sample set, namely the *Cetobacterium sp000799075* MAG. Only SNVs with more than 10X coverage across 90% of the samples were included. The results indicate that the depth of variability is mainly driven by the combination of coverage of the *Cetobacterium sp000799075* MAG and amount of host DNA among sample types. That is, the lower the host DNA content and higher the Cetobacterium coverage, the more variation is recovered for each SNVs indicating that too low coverage of MAGs leads to loss of information at the intraspecific microbial population level.

**Figure 6:**
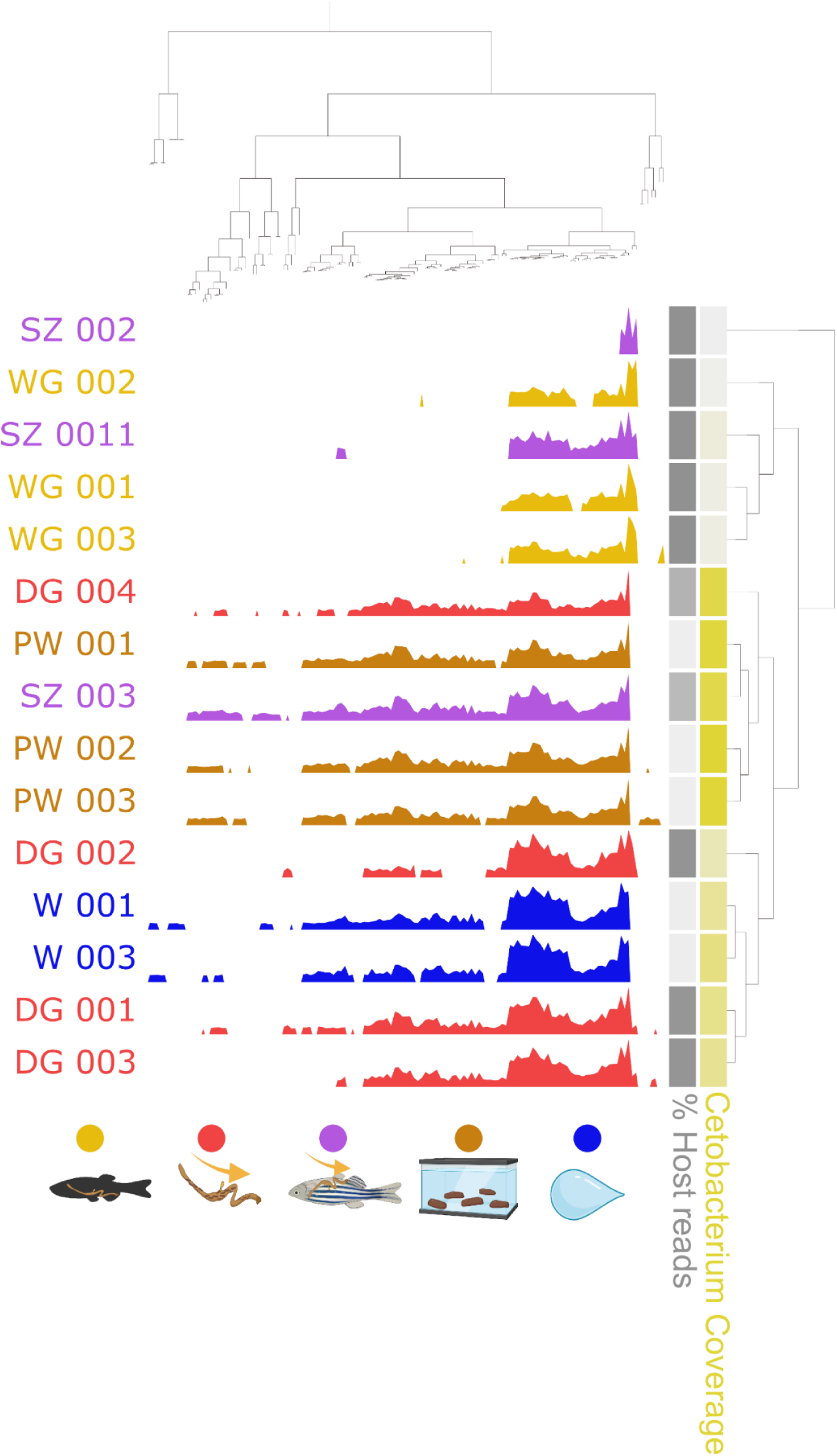
The figure shows the variability of single nucleotide variants(SNVs) in the high abundance Cetobacterium MAG. SNVs with a minimum coverage of 10X across 90% of the samples were taken into account. The variability of SNVs is hierarchically clustered using ward ordination and euclidean distance based on the similarity of variability. The y axis is similarly clustered. The layers on the right represent a gradient of Cetobacterium coverage in each sample (yellow) and a gradient of percentage of host reads in each sample.

### The representation of MAG catalogues differ between sample types

We also wanted to investigate how the data generated for this study compared to previously generated zebrafish metagenomic data. We mapped our metagenomic reads to a previously curated MAG catalogue (Gurbich et al., 2023; Richardson et al., 2023) consisting of 101 MAGs. We compared the fraction of metagenomic reads mapping to the Zebrafish Fecal v1.0 MAG catalogue to the percentage mapping to our MAG catalogue which should estimate both the representativeness of both catalogues in regards to our data and sample types (Fig. 7). The overall trend among all sample types was that a higher percentage of reads mapped to the MAG catalogue generated for this study compared to the reference Zebrafish Fecal v1.0 MAG catalogue, with the difference ranging between 5%-24%. Notably the highest fraction of reads mapped to both catalogues was observed in the intestinal content samples followed by fecal samples, squeezed gut samples, whole gut samples and finally the water samples.

**Figure 7:**
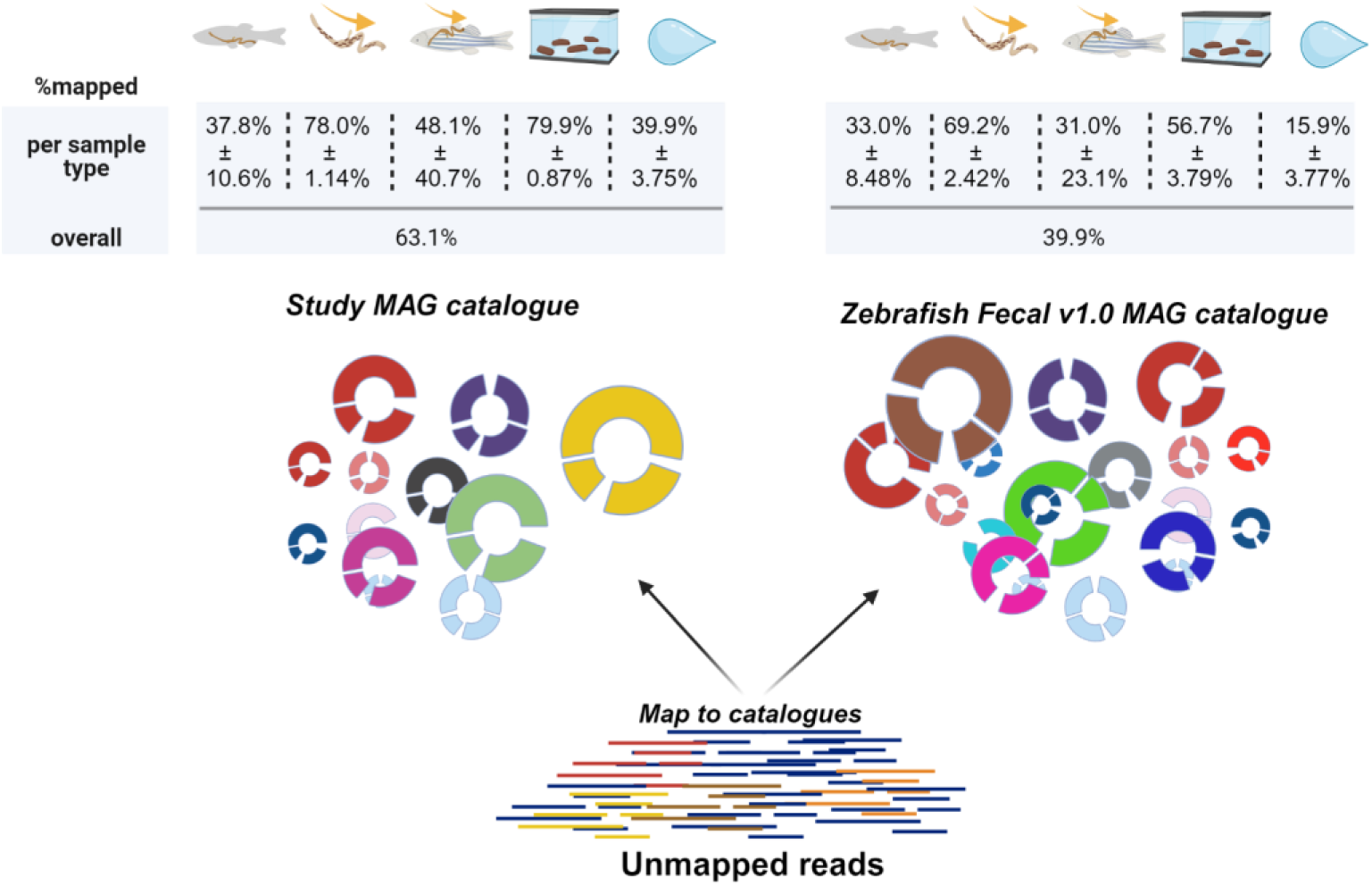
The figure shows the comparison of unmapped reads generated in this study mapped to the non-redundant MAG catalogue generated for this study and the Zebrafish Fecal v1.0 MAG catalogue from (Gurbich et al., 2023) which was generated by publicly available zebrafish shotgun metagenomic sequences. Figure created using BioRender.com

## DISCUSSION

We aimed to illustrate the impact of sampling technique on bioinformatic processing and analyses of the zebrafish gut microbiota through a typical genome resolved metagenomic pipeline. We compared four sampling techniques and included water samples as an environmental control to assess potential environmental contamination from water related microbes. We compared the sample types on practicality and consistency of sampling, through quality filtering, assembly, MAG recovery, microbiota composition and more.

Considering a sample that constitutes a good representative for the zebrafish gut microbiota, sampling the whole gut seems like a good idea at first glance since in theory it should capture the entirety of what is in the gut and even potential intracellular gut bacteria without potential contamination from the environment. However, considering the results, it may not be the best representative sample. The sheer amount of host reads (99%) leads to low number of microbial reads which leads to a whole host of problems with downstream analyses. Even though we sequenced 20G per sample the sequencing depth of the microbial content of the whole gut samples was insufficient (Fig.3A). We only recovered two MAGs, which is low compared with the rest of the sample types and what we expected from existing literature (Kayani et al., 2021, 2022). Those MAGs, belonging to the genera of *Cetobacterium* and *Aeromonas,* were however the ones that dominated the relative abundance across the sample set. This may indicate that despite the low metagenomic content in the whole gut sample, the potential for monitoring the dominating bacteria in the gut is there. We encountered similar problems with sampling the intestinal content, although to a lesser extent.

The content of the zebrafish intestinal tract should, like the whole gut samples, be a good representative of the zebrafish gut microbiota. Although more time consuming in time spent per sample, the intestinal content had lower amount of host DNA than the whole gut samples. Although 87% host DNA is a large fraction it is an improvement from the whole gut samples. Rarefaction analyses indicated sufficient sequencing depth and saturated curves although the diversity seems lesser than compared with the fecal samples. Despite the overall higher quality of data compared to the whole gut samples the microbiota profiles largely reflect a similar composition to the whole gut samples, if the non-redundant MAG catalogue is followed, with Cetobacteirum and Aeromonas dominating the composition. Overall the intestinal content samples seem to provide a microbiota resolution that is an intermediate between whole gut samples and fecal samples.

We included squeezed gut samples as we thought they would be a potentially good alternative to fecal samples. That is, sampling the feces from the fish without dissecting out the gut and without the feces coming into contact with the surrounding water, thus largely eliminating the potential problem for host and environmental contamination. However, according to the results, unfortunately, the sampling was inconsistent. Analyses of two of the three samples taken indicated insufficient sequencing depth and a similar host content as in the whole gut samples, in fact they largely reflected the whole gut samples throughout the bioinformatic processing and analyses. However, a single sample only had 55% host content, indicating sufficient sequencing depth and larger diversity compared to the intestinal content samples. This single sample is thus likely largely responsible for the 16 MAGs that we recovered from the binning efforts of the assembled contigs for the squeezed gut samples.

We believe that if this sampling method can be improved in terms of consistency, it lessens the risk associated with potential environmental contamination as in the feces sampled directly from the tank water. Interestingly the single sample higher in diversity largely reflected the composition and diversity of the fecal samples, and a microbial fraction largely absent in the intestinal content and whole gut samples.

Sampling feces directly from the tank is the most prevalent method that has been used in the existing literature of shotgun metagenomic sequencing of the zebrafish microbiota (Gaulke et al., 2020; Kayani et al., 2021, 2022). The sampling effort indicated very low host contamination (<1%), sufficient sequencing depth and recovery of 23 MAGs, moreover the three replicate samples were largely consistent throughout the analyses. The fecal samples seem to reveal a microbial fraction not present in the whole gut or intestinal content samples. In this study the fecal samples were not from individual fish but rather represent a group level within each tank. For obtaining an individual resolution, (Gaulke et al., 2019) provide a solution where individual fish were placed in individual tanks overnight and fecal matter sampled. Whether such a method produces an additional discernible tank effect should be investigated. The fecal samples raised two questions, one regarding the composition of the microbiota, as it was discernible different from the other samples and secondly regarding environmental contamination.

Starting with the overall microbiota composition it is clear that two MAGs dominate the microbiota composition (the *Cetobacterium sp000799075* MAG and *Aeromonas* MAG), These have been found in high abundance in both the previous shotgun metagenomic studies and in 16S based studies. This study may simply reflect the microbiota of the zebrafish in the facilities in which the study was conducted, as facility/study specific microbiota are well documented (Roeselers et al., 2011; Sharpton et al., 2021). What is of bigger interest is the difference in composition and diversity between the intestinal content samples and the fecal samples. This might simply reflect the added microbial fraction not captured by the intestinal content samples. It could also be that the fecal samples include microbes that are simply passing through. Otherwise, it may just be that the fecal samples and the intestinal content samples just represent different “niches” or different parts of the zebrafish gut microbiota. The number of MAGs generated in this study is perhaps on first glance rather low, but considering the high host content and the abundance of the most abundant MAGs this is perhaps to be expected. However, future studies are expected to have larger sample sizes compared to this pilot, and should improve the binning efforts and yield a higher number of MAGs.

One of the main concerns and basis of the study was the potential contamination of fecal samples from the surrounding water. We therefore attempted to apply a functional approach to the estimation of the extent of contamination. In terms of functional abundance, the results indicated a low amount of contamination in the fecal samples. The results also indicate that the water samples are more contaminated from the fecal samples than the fecal samples are potentially contaminated by the water samples. This may be the case due to a higher biomass in the fecal samples, potentially overwhelming the water microbiota community. One might also assume that MAGs or genes with higher coverage in the water samples should originate from the water, however one can never exclude the possibility that a high coverage MAG from the water lives in a low abundance niche in the zebrafish gut. Therefore it is difficult to say whether actual water contamination occurs in the fecal samples, or if it is simply something that is present in low abundance in the fecal samples. Furthermore, it is difficult to say whether increased sample numbers and sequencing effort would potentiate or reduce this contamination risk. These results are based on coverage of genes and thus any analytical approach using presence/absence of genes or metabolic pathways need to be further scrutinised as a large fraction of gene calls is shared between the environmental water microbiota (Fig.5A) and the intestinal microbiota representative sample types. Despite the low potential water contamination of the fecal samples, taken together the results indicate that caution should be taken when interpreting results from fecal matter sampled directly from the tank, and that water samples serve as an important environmental control that can help remove MAGs more likely to be sourced from the water.

We further compared our non-redundant MAG catalogue with a previously published MAG Zebrafish fecal catalogue (Gurbich et al., 2023) consisting of 101 MAGs. Overall the catalogue produced in this study MAGs performed better compared to the other catalogue. This is not surprising as the catalogue was generated from our data and should thus suit it better. However the difference between the percentages of reads mapping back to the catalogues was smaller than expected. This demonstrates the usefulness of using other reference MAG catalogues, as having a comprehensive MAG catalogue can make data processing and analyses more straightforward. One may for example capture a higher fraction of reads for downstream analyses otherwise lost in the local assembly process where rare species may not be well represented. This is particularly relevant to zebrafish considering the facility and study related differences documented in the 16S zebrafish microbiota literature (Roeselers et al., 2011; Sharpton et al., 2021).

The number of MAGs recovered per sample type is relatively low compared to existing literature in zebrafish (Gaulke et al., 2020; Gurbich et al., 2023; Kayani et al., 2021, 2022). However, considering the limited per-group sample number it is expected that the number of MAGs will increase with increased sample size due to increased binning power. The small per group sample size – ranging from two to four samples per group (only 15 samples in total) led us to largely compare the sample types qualitatively. We decided on a small-sample-size-deeper-sequencing instead of larger-sample-size-shallower-sequencing approach since we expected a high host DNA content in most of the sample types, based on previously published literature of fish microbiota studies (Collins et al., 2021; Hennersdorf et al., 2016; Rasmussen et al., 2021; Riiser et al., 2019).

The amount of host DNA is clearly different between the sample types, resulting in lower resolution of the microbial diversity. However this is not necessarily detrimental. If there is combined interest in the host genome and microbiota, sequencing the whole gut or the squeezed gut may be a good option to retrieve whole host genomes along with the microbial fraction in one sequencing run (Marcos et al., 2022).

For studies simply aiming to study the zebrafish microbiota composition and function, the amount of host DNA in the host derived samples is an obvious problem. A simple approach to increase the resolution of the microbiota is to increase the sequencing depth. However this is poorly scalable, in terms of cost, sample number, data produced and computational resources. Other methods to reduce host contamination such as chemical host depletion pre or post DNA extraction, microbial enrichment and host depletion by adaptive sequencing have been applied and compared to varying degrees (Horz et al., 2008; Marotz et al., 2018; Marquet et al., 2022; Yap et al., 2020). These promising alternatives to deep-sequencing approaches are interesting tools to apply to intestinal zebrafish samples for shotgun or long-read metagenomic sequencing. As this study simply aimed to compare and evaluate sample types these approaches were not tested. Overall this study indicates that the choice of sample type matters and underscores the importance of environmental controls, which allows for more scrutiny in analyses. We have attempted to summarise the results from the study (Table 2) and hope that it can be of use to researchers planning to undertake a genome-resolved shotgun metagenomic approach of the zebrafish microbiota.

**TABLE 2:**
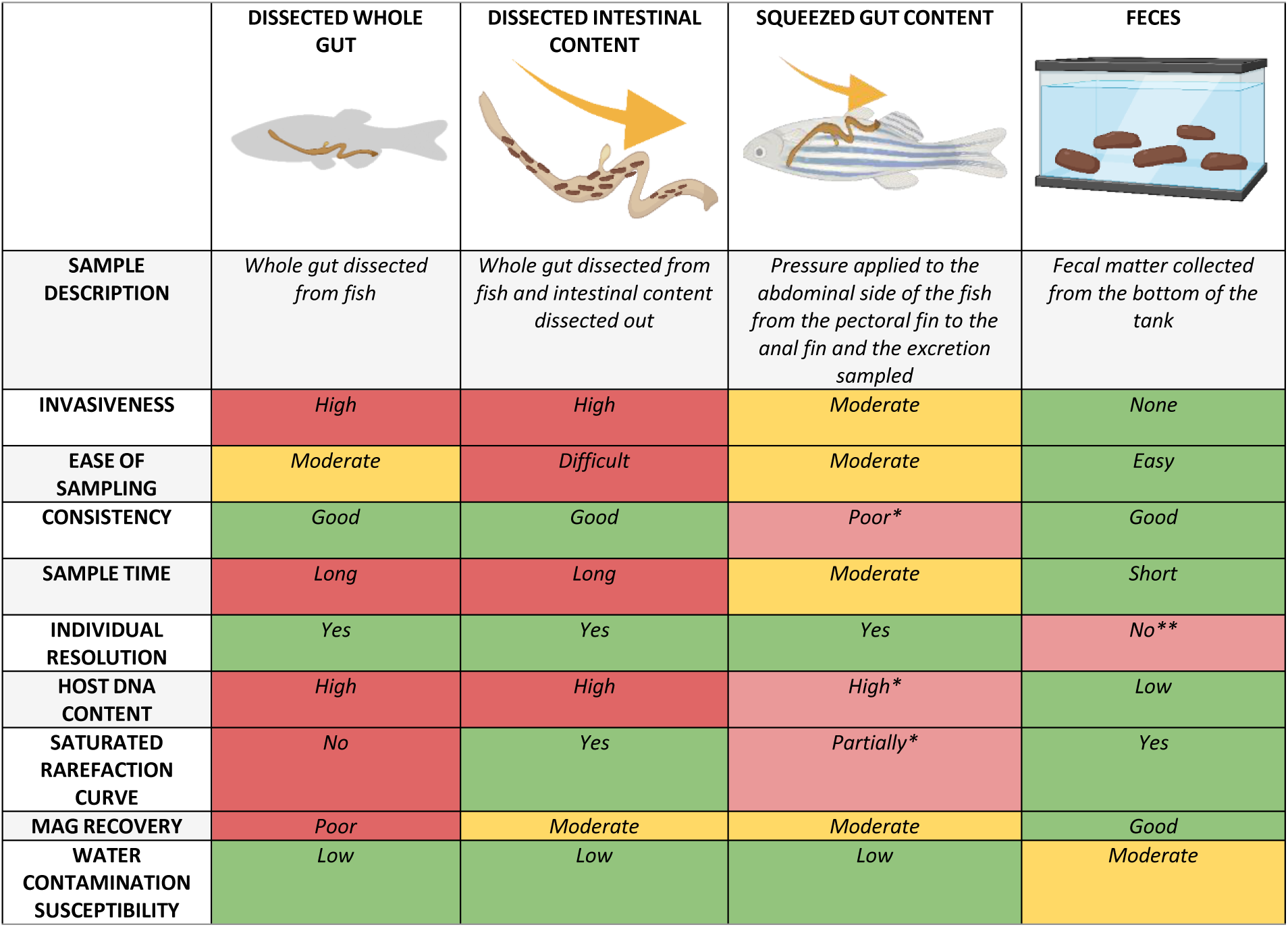
The table summarises the main results. *Potential for optimization and improvement in sampling strategy. **Can be used for individual resolution as described by (Gaulke et al., 2019)

|  | DISSECTED WHOLE GUT | DISSECTED INTESTINAL CONTENT | SQUEEZED GUT CONTENT | FECES |
| --- | --- | --- | --- | --- |
| SAMPLE DESCRIPTION | Whole gut dissected from fish | Whole gut dissected from fish and intestinal content dissected out | Pressure applied to the abdominal side of the fish from the pectoral fin to the anal fin and the excretion sampled | Fecal matter collected from the bottom of the tank |
| INVASIVENESS | High | High | Moderate | None |
| EASE OF SAMPLING | Moderate | Difficult | Moderate | Easy |
| CONSISTENCY | Good | Good | Poor* | Good |
| SAMPLE TIME | Long | Long | Moderate | Short |
| INDIVIDUAL RESOLUTION | Yes | Yes | Yes | No** |
| HOST DNA CONTENT | High | High | High* | Low |
| SATURATED RAREFACTION CURVE | No | Yes | Partially* | Yes |
| MAG RECOVERY | Poor | Moderate | Moderate | Good |
| WATER CONTAMINATION SUSCEPTIBILITY | Low | Low | Low | Moderate |

## DATA AVAILABILITY

Associated data, raw sequence reads and MAGs will be publically available upon publication

## ACKNOWLEDGEMENTS

This project was funded by The Danish National Research Foundation award (CEH – DNRF143) and a Carlsberg Foundation award to M.T.L. (CF21-0356). We would like to thank Sara Vebæk Gelskov and Carlota Marola Fernandez Gonzales for assistance with zebrafish husbandry. We would further like to thank Tom O. Delmont at Genoscope & CEH, and Jacob Agerbo Rasmussen for their expert input in analyses.

## DECLARATION OF INTERESTS

None declared

## AUTHOR CONTRIBUTION

M.T.L., E.A.T. and S.B.H. conceived the study with input from L.V.G.J. Sampling was organised and performed by L.V.G.J., M.T.L, E.A.T. and S.B.H. E.A.T. and S.B.H carried out the laboratory work and computational analyses. E.A.T. wrote the paper with input from all authors.

## Notes

### Competing Interest Statement

The authors have declared no competing interest.

