## Supplementary file S2 for "Sampling the Zebrafish gut Microbiota – A Genome Resolved Metagenomic Approach"

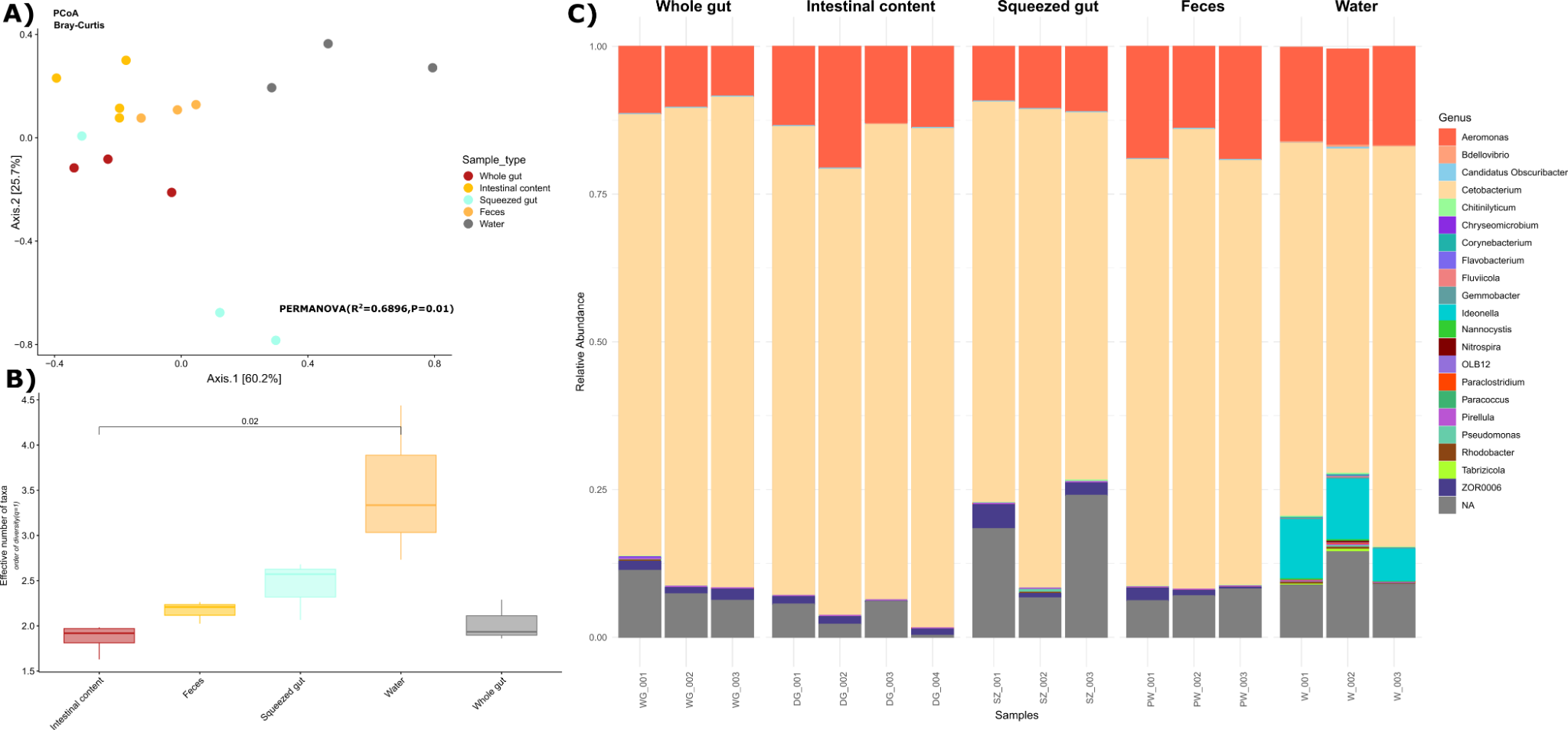

**Figure S1: 16S analyses and comparison among sample types largely reflects the shotgun sequencing dataset.**

*The figure depicts a summary of analyses of the V3-V4 16SrRNA gene region of the samples taken. A) A PCoA using Bray-Curtis distance of the samples, the dots are coloured according to sample type. B) Alpha diversity calculated using Hill numbers in order of diversity q=1. The x-axis and the colour of the boxes differentiate between the sample types. The plot indicates an increase in diversity in the following order from lowest to highest: Whole gut, Intersintal content, Feces, squeezed gut and Water samples. C) A stacked barplot depicting the relative abundance of Genera (coloured) among the different sample types.*

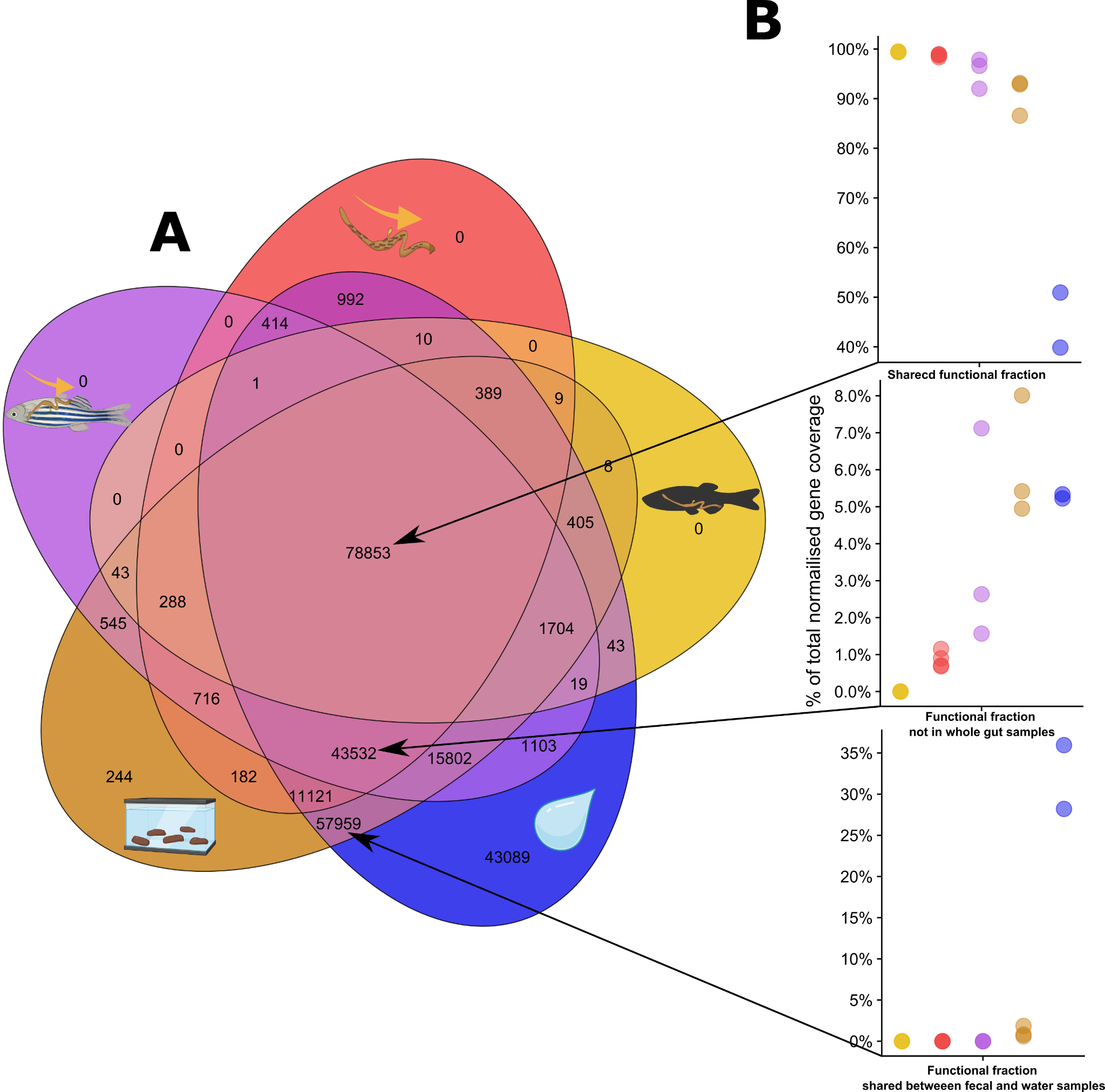

**Figure S2: Comparative analyses of genes assigned Pfam annotations and their functional abundance among sample types**

*A) Venn diagram of number of gene calls with assigned Pfam categories across between sample types. B) percentage of normalised gene coverage of genes in fractions of the venn diagram, as indicated by the arrows, among sample types. The colors and icons represent the different sample types. The colours and icons indicate the sampling type, yellow is whole gut samples, red is Intestinal content samples, purple is squeezed gut samples, brown is feces and blue is water samples.*
